# Exploring Nonlinear Dynamics In Brain Functionality Through Phase Portraits And Fuzzy Recurrence Plots

**DOI:** 10.1101/2023.07.06.547922

**Authors:** Qiang Li, Vince D Calhoun, Tuan D. Pham, Armin Iraji

**Affiliations:** Tri-Institutional Center for Translational Research in Neuroimaging and Data Science (TReNDS), Georgia State University, Georgia Institute of Technology, and Emory University, Atlanta, GA, 30303, USA; Barts and The London School of Medicine and Dentistry, Queen Mary University of London, Turner Street, London E1 2AD, UK

**Keywords:** Nonlinear Dynamics, Biophysics, Phase Portrait, Fuzzy Recurrence Plots, fMRI, Functional Connectivity Descriptors

## Abstract

Much of the complexity and diversity found in nature is driven by nonlinear phenomena, and this holds true for the brain. Nonlinear dynamics theory has been successfully utilized in explaining brain functions from a biophysics standpoint, and the field of statistical physics continues to make substantial progress in understanding brain connectivity and function. This study delves into complex brain functional connectivity using biophysical nonlinear dynamics approaches. We aim to uncover hidden information in high-dimensional and nonlinear neural signals, with the hope of providing a useful tool for analyzing information transitions in functionally complex networks. By utilizing phase portraits and fuzzy recurrence plots, we investigated the latent information in the functional connectivity of complex brain networks. Our numerical experiments, which include synthetic linear dynamics neural time series and a biophysically realistic neural mass model, showed that phase portraits and fuzzy recurrence plots are highly sensitive to changes in neural dynamics and can also be used to predict functional connectivity based on structural connectivity. Furthermore, the results showed that phase trajectories of neuronal activity encode low-dimensional dynamics, and the geometric properties of the limit-cycle attractor formed by the phase portraits can be used to explain the neurodynamics. Additionally, our results showed that the phase portrait and fuzzy recurrence plots can be used as functional connectivity descriptors, and both metrics were able to capture and explain nonlinear dynamics behavior during specific cognitive tasks. In conclusion, our findings suggest that phase portraits and fuzzy recurrence plots could be highly effective as functional connectivity descriptors, providing valuable insights into nonlinear dynamics in the brain.

**Significance:** Here we report that phase trajectories and fuzzy recurrence plots can serve as descriptors of nonlinear functional dynamics networks. Both metrics are highly sensitive to variations in neural signals and are powerful tools for capturing distinct patterns from brain signals, making brain fingerprinting possible. This has significant implications for understanding brain state dynamics through phase trajectories and fuzzy recurrence plots, as both metrics are effective in identifying nonlinear dynamics patterns.

## 1 Introduction

The complexity of brain activity is represented by the nonlinearity of neural signals [1–4]. The brain neurons fire in nonlinear patterns, meaning that even when they are connected, their firing rates may differ [5, 6]. Nonlinear patterns are often associated with more complex cognitive processes, making them crucial to understanding how brain networks process information.

Considering that nonlinear dynamics and the synchronization of large-scale neural networks in the brain are closely interconnected [7, 8], applying nonlinear dynamical approaches to study complex interactions between brain regions is essential for understanding information processing in the human brain. In addition, examining the interaction between different brain regions can provide insight into the role of large-scale neuronal synchronization in cognitive processes. There may be a strong relationship between changes in functional connectivity and variations in neural dynamics [9–11].

Functional magnetic resonance imaging (fMRI) measures functional connectivity, revealing the statistical coupling between different brain regions during resting-state or specific task conditions [12–19]. It also helps identify irregularities in connectivity that may be related to specific neurological conditions. Quantifying functional connectivity can be achieved using a variety of approaches. These methods include linear and nonlinear tools from statistics, data analysis, and time series analysis, as well as techniques from differential equations, dynamical systems, and bifurcation theory [20]. Specific methods encompass Granger causal connectivity analysis [21], phase synchronization connectivity analysis [22], independent component analysis [23, 24], partial correlation [25], mutual information [26], manifold learning algorithms [27–29], and diffusion maps [20]. Additionally, approaches such as second-order statistical analysis [30], wavelets [31, 32], artificial neural network analysis [33, 34], higher-order information interaction metrics [35–40], and advanced source modeling [41–44] are also utilized. All of these methodologies are valuable for investigating cognitive processes and neurological conditions [45–48]. However, while some methods focus on capturing nonlinear relationships among components [49, 50], none specifically target the high-dimensional nonlinear dynamics of brain network behavior.

To more directly reveal the nonlinear dynamics and behavior of systems, as well as the dynamic information hidden within time series, metrics such as phase trajectories and fuzzy recurrence plots (FRP) have been successfully applied to various time series analyses [51]. These methods are increasingly used to analyze the nonlinear dynamics of neural activity [52–55]. Phase trajectories track changes in neural activity over time, while FRP is based on the idea that fuzzy logic can represent signals from different brain regions, aiding in the detection of previously unrecognized complex patterns [51]. Consequently, phase trajectories and FRP have been utilized in several sub-fields, including neurophysiology [53, 54, 56]. They have also been applied to investigate neural networks, monitor changes in brain activity over time, and explore the causes of various neurological diseases [57].

In this work, we explore the neural nonlinear dynamics of complex brain networks using phase trajectories and FRP from a biophysics perspective. We demonstrate that phase trajectories and FRP are sensitive to the neural variability within complex brain networks through both numerical simulations and real brain developmental fMRI experiments, showing their potential for capturing brain fingerprinting properties. In our numerical experiments, we generate neural time series from both physically and biophysically realistic models, and we illustrate how phase trajectories and FRP can effectively capture the nonlinear dynamics of neural signals. Additionally, we apply these metrics to a real fMRI dataset to examine how brain networks evolve with age, highlighting the complex functional transformations that occur. Estimating the changes in coupled information during brain development in both children and adults provides valuable insights into the evolution of brain network functions. Overall, our findings indicate that phase trajectories and FRP are effective tools for identifying the complex nonlinear dynamics of brain networks.

The remainder of this paper is organized as follows: Section 2 introduces the methodologies and results, demonstrating that phase portraits and FRP are highly sensitive to the variability of neural signals and effective in capturing the nonlinear dynamics of synthetic neural time series. Additionally, we use these metrics to estimate brain developmental connectomes from fMRI datasets, covering various brain regions, and present some intriguing findings. Sections 3 and 4 provide a general discussion and conclusion, respectively.

## 2 Methodologies and Results

### 2.1 Phase Trajectory Reconstruction

In the context of a complex nonlinear dynamic system, the determination of the state variables, which constitute the minimum set of variables required to fully describe the system’s state, is not readily available and must be inferred from observational data. This process is commonly referred to as state space reconstruction. The established approach to this problem involves the use of delay-coordinate embedding, or Taken’s delay embedding, as described in the literature [58].

This involves the construction of a new vector in a reconstructed space using a series of previous measurements of a single scalar variable *x* of the dynamic system. The resulting *d*-dimensional vector 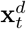 is formed from *d* time-delayed measurements of *x*_*t*_, according to the following procedure:

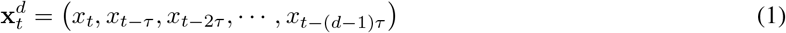

Many strategies exist to obtain the optimal embedding delay *τ* and the dimension parameter *d* when utilizing the delay-coordinate embedding method. In particular, autocorrelation and mutual information approaches have been used to estimate time delay *τ* [59, 60], while Cao’s methodology is a well-liked option for calculating the embedding dimension *d* [61]. Significantly, the values of *d* and *τ* can have a substantial effect on the precision of phase space estimates. In particular, a low value of *d* may not adequately depict the underlying dynamics of the system, whereas an excessively large number may result in computational needs that are unrealistic. In actual applications, such as functional brain network analysis for unknown systems, it can be difficult to generate accurate estimates of *τ* and *d* using autocorrelation or the Cao approach alone. In such situations, an iterative search may be required to discover the ideal values of these parameters based on early estimates.

### 2.2 Fuzzy Recurrence Plots

A fuzzy recurrence plot (FRP) is an extension of recurrence plots (RP). The RP^1^ is a sophisticated way to visualize multivariate nonlinear data [51] (here refers to neural signals). In essence, this is a graph representing a binary matrix, with members indicating recurrences of data states or phases in specific time points. Recurrence is a fundamental characteristic of deterministic dynamical systems, like nonlinear or chaotic systems, e.g., brain. Higher dimensional neural signal can only be viewed by projection into 2D or 3D sub-spaces via principal component analysis and manifold learning [27–29]. However, when compared to recurrence plots, some buried information is lost, and information transitory in phase domain with time is inadequately preserved. The RP can be mathematically expressed as follows:

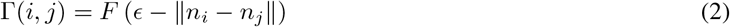

where *ϵ* indicates a threshold for defining the similarity or dissimilarity of the pair (*n*_*i*_, *n*_*j*_). *F* refers to a unit step function for producing 0 or 1 if (*ϵ −* |*n*_*i*_ − *n*_*j*_|) *<* 0 or otherwise, respectively. A RP is a binary symmetrical matrix, according to Equation 2. It displays pairs of time points at which the trajectory is at the same spot (represented by a black dot) or is not (represented by a white dot).

To address the challenge of establishing a cutoff point for similarity and the binary limitations of expressing recurrence using the RP method, a FRP was developed as an enhanced approach [51]. Given an embedding dimension *m* and a time delay *τ*, the phase-space reconstruction of the original time series (*t*_1_, *t*_2_, …, *t*_*M*_) results in **X** = (**x**_1_, …, **x**_*N*_), where *N* = *M* − (*m* − 1)*τ*. The reconstructed elements of **X** can be obtained as [58],

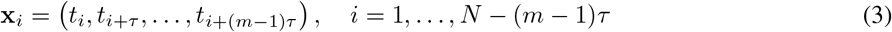

Using the reconstructed phase space **X**, a FRP, denoted as **Π**, can be created as a grayscale image to illustrate the recurrence of a dynamical system. Specifically, a FRP is a square matrix of fuzzy membership grades that quantify the similarity between each pair of points in the reconstructed phase space trajectory of the system. In phase space, the FRP illustrates the trajectory’s recurrence as a grayscale image with values ranging from [0, 1], where 0 represents black, 1 represents white, and intermediate values correspond to shades of gray. It can be mathematically expressed as follows:

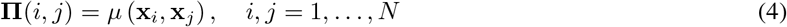

where *µ* (**x**_*i*_, **x**_*j*_) ∈ [0, 1] represents the fuzzy membership of similarity between **x**_*i*_ and **x**_*j*_. The FRP method applies the following properties:

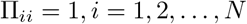

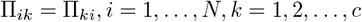

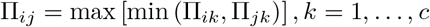

- Reflexivity
- Symmetry
- Transitivity

where Π_*ik*_ = *µ* (**x**_*i*_, **v**_*k*_) can be optimally determined using the fuzzy c-means (FCM) algorithm [62]. The FCM algorithm is employed to create the FRP by partitioning **X** into *c* clusters. This algorithm assigns a fuzzy membership grade, denoted as Π_*ik*_, which takes values in [0, 1], to each **x**_*i*_ (where *i* = 1, 2, …, *N*) in relation to each cluster center **v**_*k*_ (where *k* = 1, 2, …, *c*). As previously mentioned, the values of Π_*ik*_ are optimally determined by the FCM algorithm, which aims to minimize the following objective function:

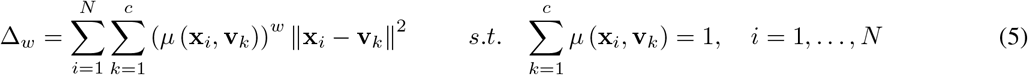

where *w* is the fuzzy weighting exponent, and we used *w* = 2 in our study. The FCM objective function is minimized through a numerical scheme that iteratively updates the fuzzy membership grades and cluster centers until the fuzzy membership grades converge (the maximum number of iterations, set to 100, and the convergence tolerance, which is 1e-5 in our study). The fuzzy membership grades and cluster centers in the FCM are updated as follows:

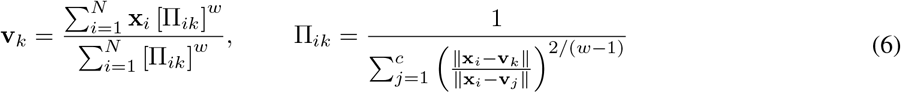

In summary, a FRP is defined as a square grayscale matrix,

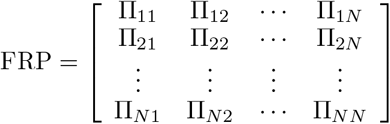

Time series are transformed into objects in a space known as dynamical system phase space, with each object in the phase space referred to as a dynamical system point. This phase space representation facilitates the discovery of underlying properties in complex data compared to the original time series [1], making it an especially valuable transformation. The embedding dimension *m* is selected to capture the primary dimensions of the neural signals, while the time delay *τ* and the number of clusters *c* in the time series are chosen to optimize the extraction of nonlinear dynamics features.

### 2.3 Numerical Experiments

We include two main sections, with the first half devoted to numerical experiments to verify the validity and sensitivity of the applied methods. In the second part, we used real brain developmental fMRI data, specifically from a public database that tracks how the brain changes over age using phase portraits and FRP.

#### 2.3.1 Recurrence Plots Capture the Variability of Time Series

Here, we construct two time series with time length *n* = 100 in Equation 7. One presents a periodic wave, while the other presents a drift scenario. The purpose of this numerical experiment is to demonstrate how sensitive the phase trajectory and RP are to the variability of time series.

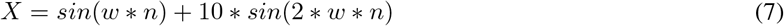

It generates a periodic neural time series for *w* = *w*_*p*_ = 2 * *π/*5 and a synthetic drift signal for *w* = *w*_*d*_ = 2 * *π/*2. Based on Equation 2, we could have a graph representing a binary matrix, with members indicating recurrences of neural states (see Figure. 1).

**Figure 1.**
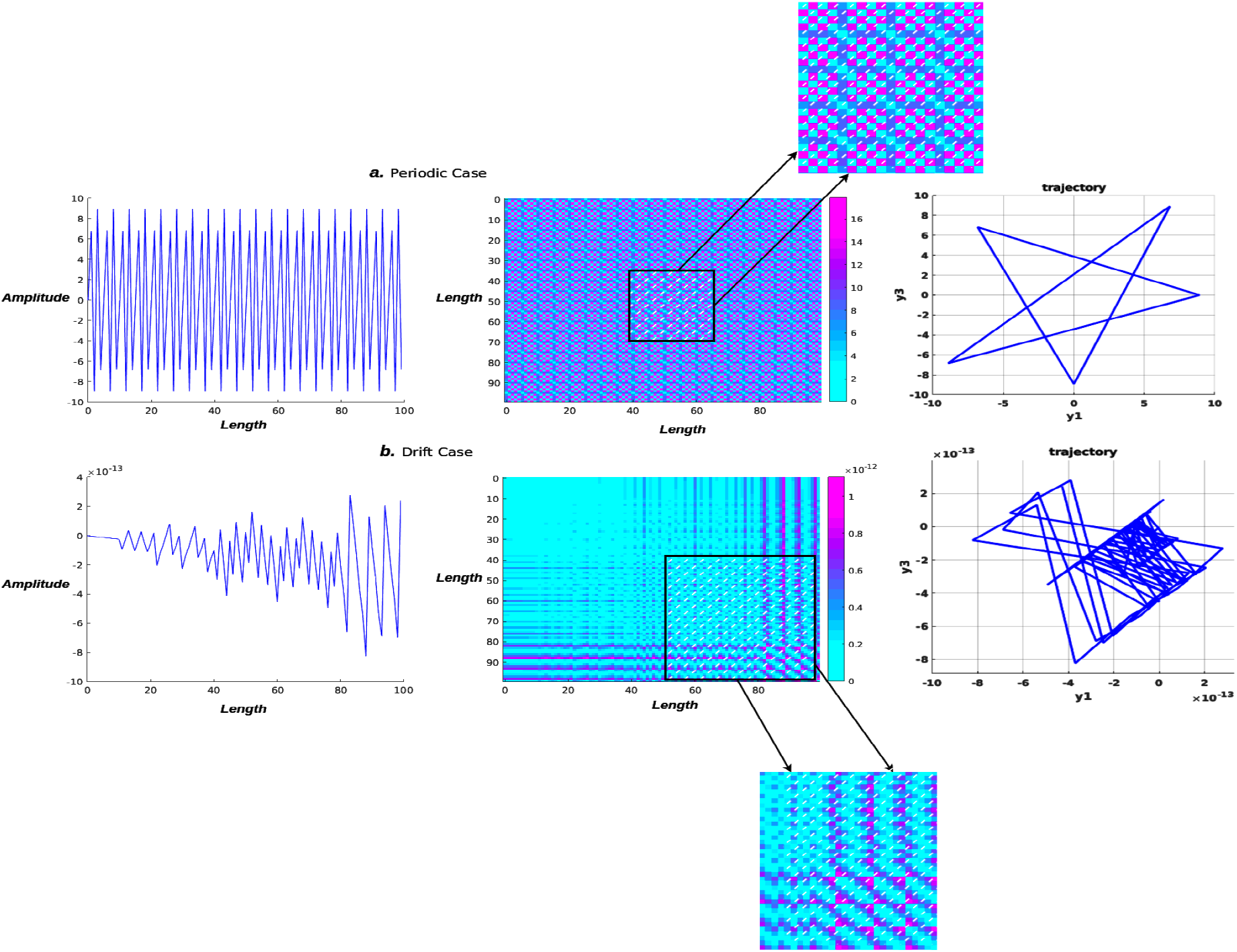
Recurrence plot and phase trajectory visualization. The periodicity of the neural signal (top row) may be seen in the recurrence plot, which shows the same pattern repeated over and over again in 2D fuzzy recurrences. The drift neural signal (second row), on the other hand, exhibits a completely distinct pattern depending on the time interval. The vertical or horizontal axis denotes a period of time during which the neural state remains constant or changes very slowly. In both cases, phase portraits are plotted, and they all show totally different dynamic trajectories.

#### 2.3.2 Fuzzy Recurrence Plots in Analyzing Linear Dynamics of Synthetic Time Series and Noise Perturbations

After demonstrating RP’s ability to detect state change sensitivity in a previous numerical experiment, we next examined the linear dynamics of synthetic neural time series and the sensitivity of FRP to noise in this simulation scenario. First, we use Equation 8 to generate a simple periodic neural signal,

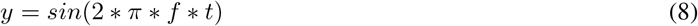

where *f* = 20 represents the frequency of the input neural signals and *t* = 1 : 100 represents the time length. In this case, the input-generated signal is shown in Figure. 2 is shown at the top left (**A**), followed by the corresponding trajectory in 3D phase space (**B**), recurrence plot (**C**), and fuzzy recurrence plot (**D**) plotted separately. Specific parameters are chosen to plot a trajectory and a fuzzy recurrence plot, such as, *m* = 3 represents the embedding dimension of a reconstructed phase space, and *τ* = 1 represents the time delay required to reconstruct phase space (see Figure. 2 top right). For the fuzzy recurrence plot, we used *c* = 8, which represents the number of clusters for fuzzy c-means clustering, and *T* = 0.1, which indicates the cutoff fuzzy membership threshold to change grayscale to black and white (see Figure. 2 bottom right).

**Figure 2.**
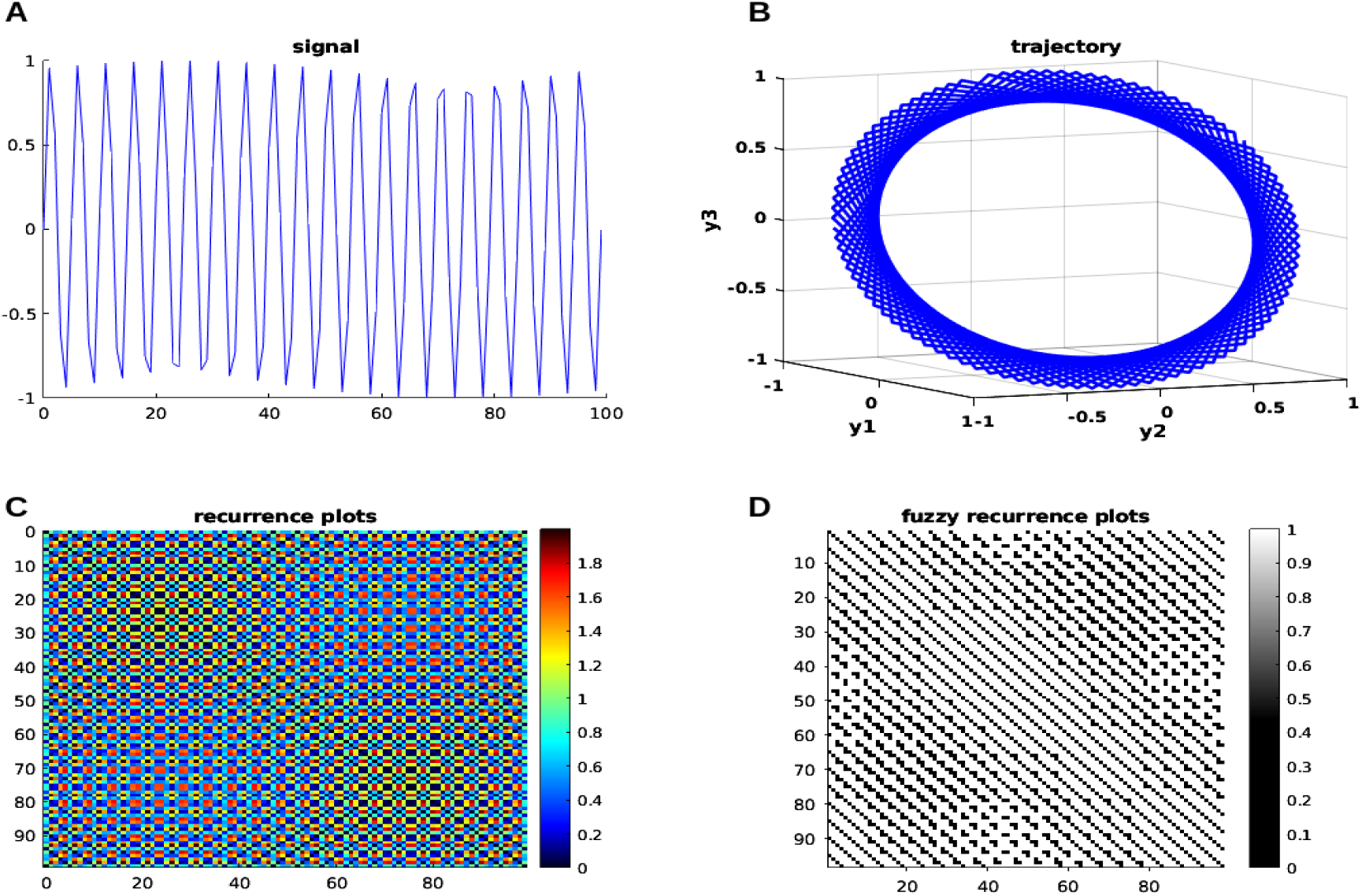
Neural modeling time series with nonlinear dynamics. The generated neural signals are on the left (**A**), and the trajectory of the input signal in 3D phase space is on the right (**B**); the recurrence plots is on the left (**C**), and the FRP is on the right (**D**). See the main text for more information.

There are several hyperparameters in FCM when estimating the FRP. To further evaluate the sensitivity of these hyperparameters, we explored the range of each parameter and summarized this information in the Table. 1.

**Table 1.** Sensitivity of the Hyperparameters in FCM.

| Hyperparameter | Value Range | Sensitivity Analysis |
| --- | --- | --- |
| Number of Clusters (c) | [2, 4, 6, 8, 10] | High Sensitivity |
| Fuzzy Weighting Exponent (w) | [1.5, 2, 2.5, 3, 3.5] | High Sensitivity |
| Maximum Iterations | [100, 200, 300, 400, 500] | Low Sensitivity |
| Convergence Tolerance | [1e-05, 5e-05, 0.0001, 0.0005, 0.001] | Moderate Sensitivity |

We found that the number of clusters (c, c=8) and the fuzzy weighting exponent (w, w=2) exhibit high sensitivity compared to the maximum number of iterations and convergence tolerance. In conclusion, we suggest that the optimal values for the number of clusters (c) and the fuzzy weighting exponent (w) depend on your specific data and should be chosen manually. However, the maximum number of iterations can typically be set to 100, and the convergence tolerance can be set to 1e-5 in general situations.

Meanwhile, we applied fuzzy Determinism (DET), fuzzy Laminarity (LAM), fuzzy Trapping Time (TT), and fuzzy Recurrence Entropy (ENT) to quantify FRP and RP [63]. These metrics were applied to three different types of time series: random noise, pink (1/f) noise, and the chaotic time series of the x-component of the Lorenz system. The results for each metric are presented in the Table. 2.

**Table 2.** Quantification Analysis of FRP and RP.

| Signal Types | DET | LAM | TT | ENT |
| --- | --- | --- | --- | --- |
| FRP ($L_{d_{\min}} = 5, L_{v_{\min}} = 5$) | | | | |
| Random Time Series | 0.6329 | 0.4915 | 1.0017 | 1.3414 |
| Pink (1/f) Noise | 0.4756 | 0.7438 | 2.4795 | 1.5631 |
| Lorenz (x-component) | 0.9938 | 0.9900 | 28.6180 | 4.3127 |
| RP ($L_{d_{\min}} = 5, L_{v_{\min}} = 5$) | | | | |
| Random Time Series | 0 | 0 | NaN | 0 |
| Pink (1/f) Noise | 0 | 0 | NaN | 0 |
| Lorenz (x-component) | 0.9982 | 0.9953 | 24.9763 | 5.6474 |

where 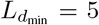 and 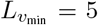 are two free parameters that refer to the minimum lengths of diagonal and vertical segments, respectively. We observe that the FRP-based and RP-based measures consistently show differences in quantifying random noise and chaotic time series (specifically, the Lorenz x-component), validating the FRP-based quantification analysis. Additionally, the FRP approach avoids the zero or NaN values that can occur with RP-based measures. Second, we found that chaotic signals exhibit significantly higher values compared to random noise signals, and both FRP and RP can effectively distinguish between chaotic and non-chaotic signals.

Now, we add white Gaussian noise to the signal with a control signal-to-noise ratio (SNR) of 12 dB in Equation 9, and the nonlinear dynamic can be presented visually in Figure. 3 to assess for a transition in the pattern.

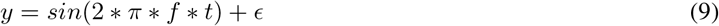

**Figure 3.**
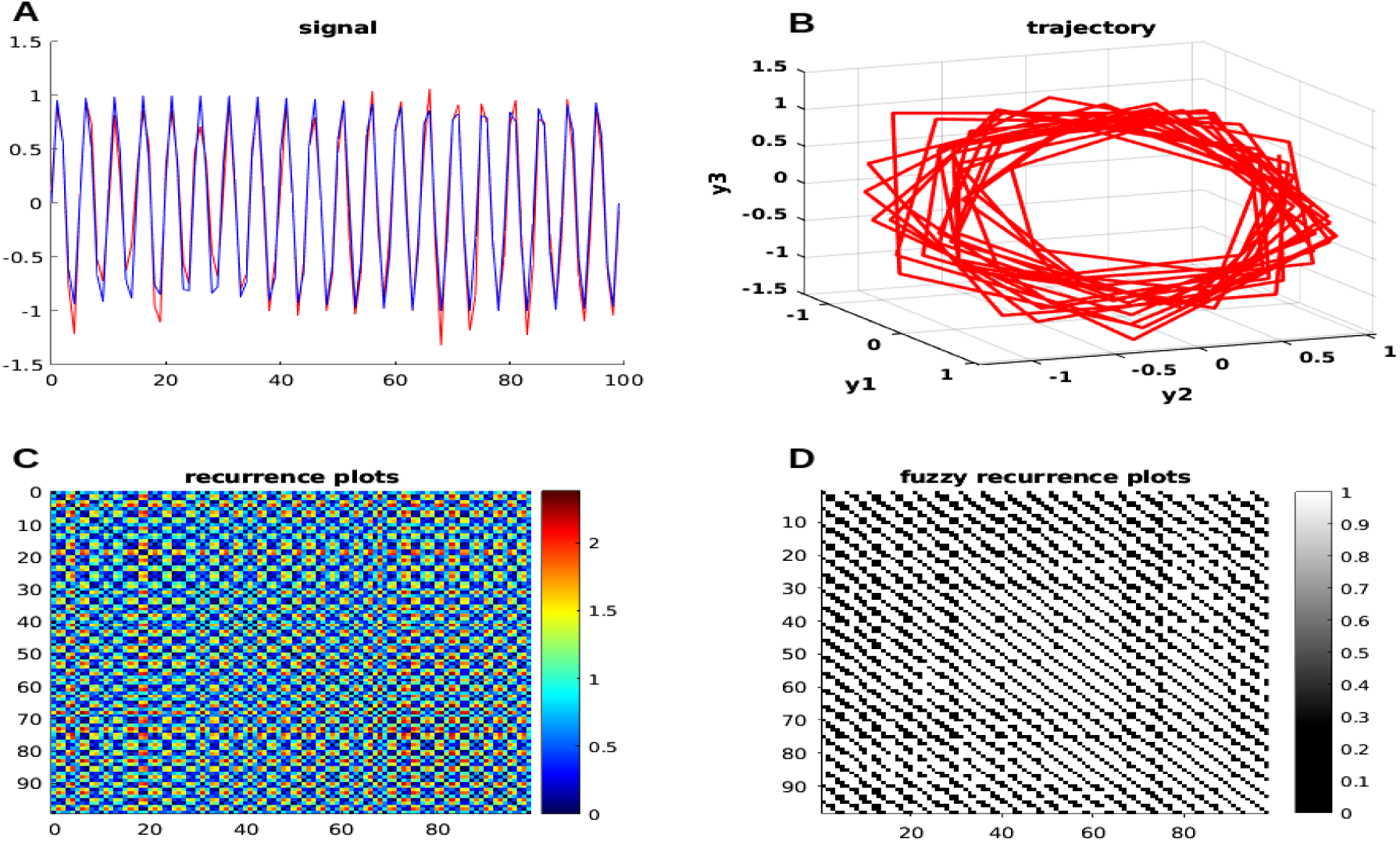
Time series with the added noise of neural simulation with nonlinear dynamics. The generated neural signals are on the left (**A**) (blue represents the original signal, while red represents the added noise signal), and the trajectory of the input signal in 3D phase space is on the right (**B**); the recurrence plots is on the left (**C**), and the FRP is on the right (**D**). See the main text for more information.

In addition, we explored different levels of SNR at 6, 12, 18, and 24 dB using FRP to assess their sensitivity to varying SNR levels. We observed that as SNR increases, all metrics, including DET, LAM, TT, and ENT, also increase, as shown in Table. 3. It reflected that FRP was sensitive to noise perturbations.

**Table 3.** Quantifying Different Levels of SNR in FRP.

| Random Time Series (SNR/dB) | DET | LAM | TT | ENT |
| --- | --- | --- | --- | --- |
| FRP ($L_{d_{\min}} = 5, L_{v_{\min}} = 5$) | | | | |
| 6 | 0.8311 | 0.4478 | 2.4161 | 2.5872 |
| 12 | 0.8555 | 0.5587 | 7.3691 | 2.9480 |
| 18 | 0.9734 | 0.9610 | 18.8726 | 4.0424 |
| 24 | 0.9960 | 0.9929 | 17.6201 | 4.7395 |

##### 2.3.3 Biophysically Realistic Whole-Brain Neural Mass Model

As we mentioned earlier, an interconnected network of brain regions makes up a model of the entire brain. Each part of the brain is represented by a neural mass model that is connected to other parts of the brain. On a mesoscopic scale, the neural mass models that represent the average activity of a brain region represent the cortical activity generated by the dynamic interplay of various neural populations subject to excitatory and inhibitory feedback [64]. For large-scale systems, like the population neural activity, a neural mass model will typically involve a set of differential equations (see Figure. 4). Therefore, these models are thought to be helpful for representing the typical activity of a sizable population of neurons, such as a region of the brain. Every simulated brain region generates its own activity, such as a population firing rate. To calibrate the model against empirical data from fMRI, the activity of each brain region can be transformed into a simulated blood-oxygenation-level-dependent (BOLD) signal, and the validity of the model can be determined by contrasting the simulated result with actual brain recordings.

**Figure 4.**
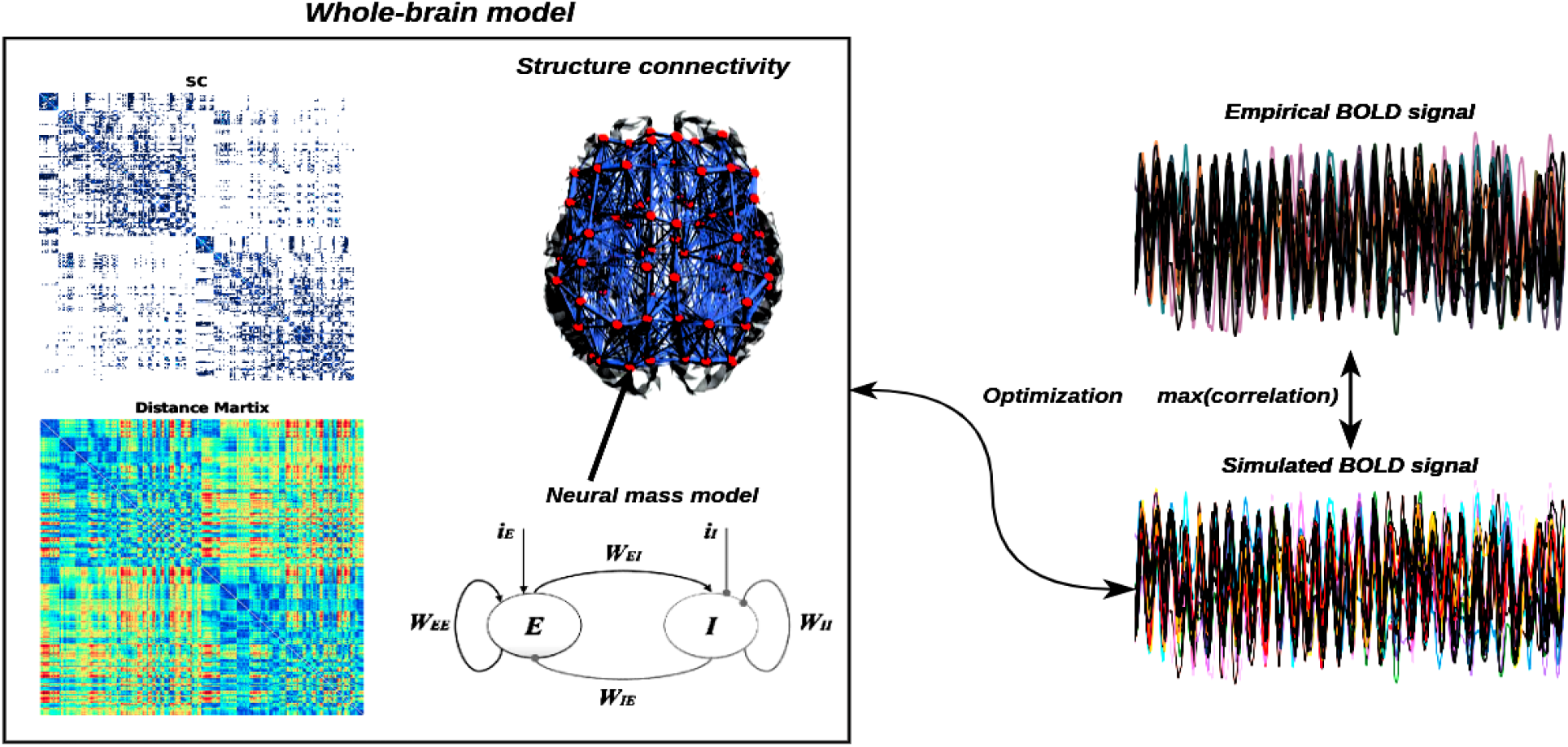
Overview of whole-brain neural mass model. The left panel shows the large-scale structure connectivity (structure connectivity coupling strength matrix and fiber distance matrix) and neural mass model considered with Excitatory(*E*)-Inhibition(*I*) couple networks (e.g., *W*_*EE*_,*W*_*II*_, *W*_*EI*_, *W*_*IE*_, and external currents input, *iE* and *iI*), and the right panel shows the simulation BOLD signal from the neural mass model, which can be optimized for maximum correlation between the empirical and simulated BOLD signal.

Our analysis is based on a system of *N* = 400 neural oscillators coupled in the connectome, considering both the connectivity strength, *C*_*ij*_, and the conduction delays between each pair of brain areas *i* and *j*. The conduction delays are defined in proportion to the fiber lengths between brain areas. Based on the Kuramoto model of coupled oscillators with time delays [65–68], so the model is defined by the following equation:

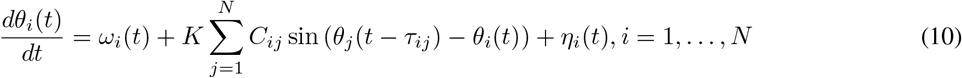

where *θ*_*i*_ is the phase of each oscillator i, *ω* = 2*πf, f* = 40 is its intrinsic frequency, the delay *τ*_*ij*_ between node *i* and node *j* is calculated using *τ*_*ij*_ = *L*_*ij*_*/V* = ⟨*τ*⟩ *L*_*ij*_*/* ⟨*L*⟩, where ⟨*L*⟩ indicates the mean fiber length, *V* represents conduction velocity, and the time delay value is 0.01 second in this study, and *η*_*i*_ is a Gaussian noise component with mean *µ* = 0 and standard deviation *σ* = 0.05. The population neural activity can be transformed into BOLD signals through the forward Balloon-Windkessel hemodynamic model [43]. Hemodynamic models, in brief, explain how changes in population neuronal activity affect the vasculature, which in turn affects blood flow, leading to alterations in vessel volume and deoxyhemoglobin content, the latter of which underlies BOLD signals. All of the parameters in the model were taken from Friston et al. [43].

Here we used the Kuramoto neural mass model to simulate the 400 brain regions of BOLD signals over 1 second (see Figure. 5**A**), the related phase trajectories are plotted in Figure. 5**B**, and their corrspording power spectrum are plotted in Figure. 5**C**. The Figure. 5**C** and **D** depict the structure connectivity and neural fiber distance matrix from Diffusion Tensor Imaging (DTI), Both sets of connectivity data were estimated from Human Connectome Project (HCP) connectivity data, and in this simulation, we used this connectivity data from Esfahlani et al [69]. The coupling between structure connectivity and functional connectivity is one of the key problems in neurophysics research because functional connectivity is constrained by structure connectivity, and the relationship between structure connectivity and functional connectivity is still unclear. However, as we mentioned before, one way to clarify this question is from the neural mass model, where we can predict and optimize predicted functional connectivity from structure connectivity based on the neural mass model. Here, we use FRP to represent predicted functional connectivity, and the means of Pearson correlation and mutual information-based functional connectivity are presented in Figure. 5**F** and **G**, respectively. Both the phase plane and the FRP can capture the nonlinear dynamics behavior in neural time series, and they can estimate predicted functional connectivity with certain accuracy, allowing us to better understand the coupling strength between structure connectivity and functional connectivity.

**Figure 5.**
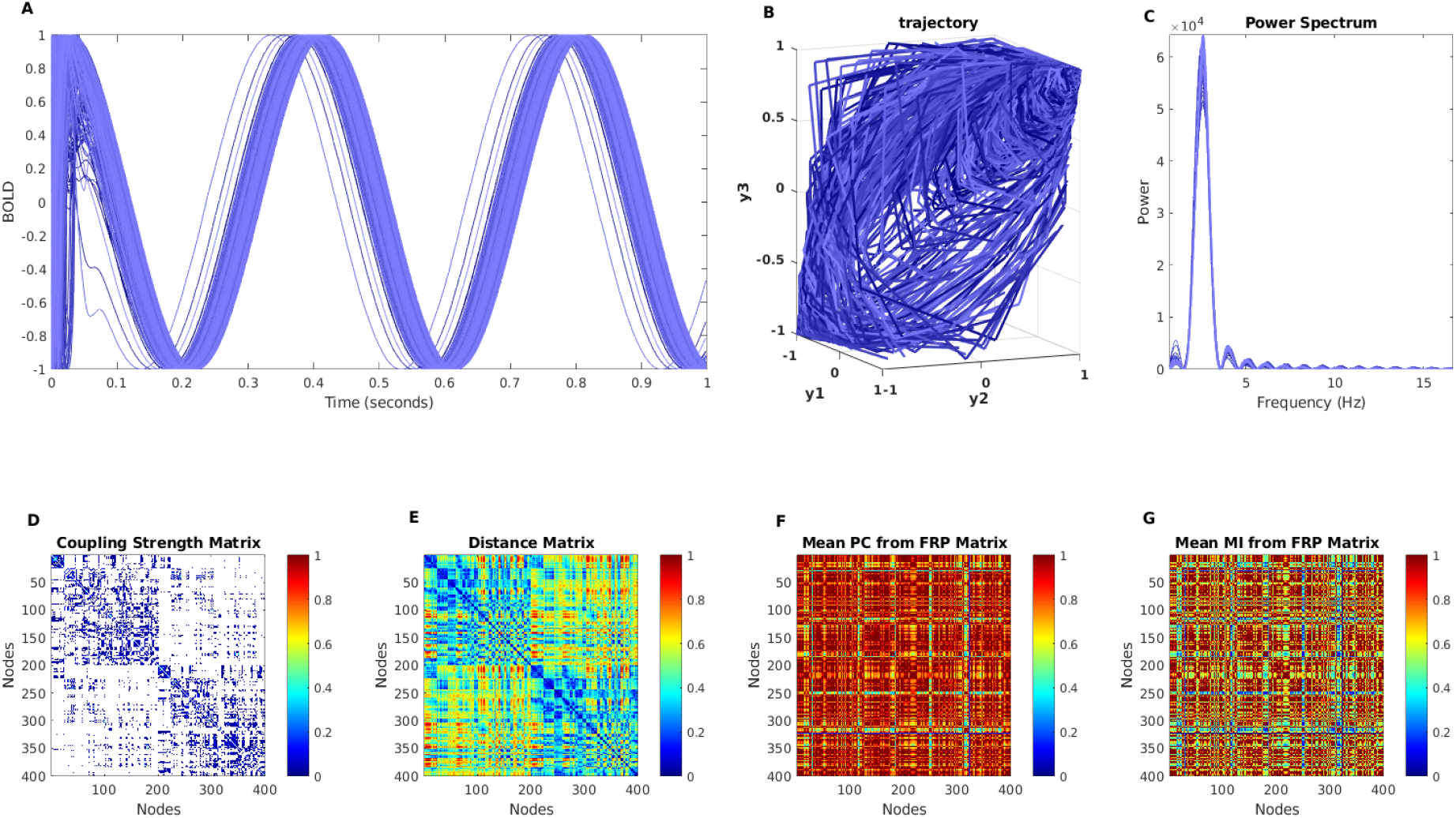
The BOLD simulation with neural mass model. The **A** shows the simulated BOLD signals from 400 brain regions with the Kuramoto model, and **B** shows the phase portrait of simulated BOLD signals with *m* = 3 and *τ* = 1. The **C** shows power spectrum corresponding to synthetic neural time series. The empirical structure connectivity (**D**), empirical fiber distance connectivity (**E**), and predicted functional connectivity(**F**,**G**) are shown in the second row, respectively.

Now by determining the global order parameter, the synchronization behavior of the dynamics system was evaluated,

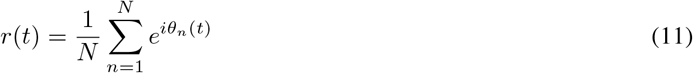

where *θ*_*n*_(*t*) is the phase of each node, and *r*(*t*) quantifies the degree of phase synchronization of the ensemble, with *r*(*t*) = 0 indicting complete asynchrony and *r*(*t*) = 1 indicating complete synchrony. In our study, we use the mean *r*(*t*) as a measure of global phase synchronization, while the standard deviation std *r*(*t*) indicates how much *r*(*t*) fluctuates in time [70, 71]. The free parameters *K* are adjusted to investigate the parameter space. To make sure our range included both weak and strong coupling, we opted to increase *K* from 0.08 to 5.

In the Kuramoto model, the free parameter, specifically coupling strength *K*, has the greatest influence on model performance. We explored the range of parameter spaces with varying coupling strengths, and meanwhile, we evaluated the order of parameters of simulated neural signals (see Figure. 6**A**). The related phase trajectories are plotted to assess the nonlinear dynamics of order parameters, and we can see that the weak coupling strength has tighter, limited cycles, while the stronger coupling strength gradually becomes shift and diffusion (see Figure. 6**B**). In Figure. 6**C** shows the corresponding power spectrum and mean of the order parameter with different coupling strengths (e.g. *K* = [0.08, 0.4, 0.6, 0.8, 1, 5]) (Figure. 6**D**). As we stated before, the global order parameter is a metric used to examine the level of neural synchrony, and we mapped the global order parameter via the *ENIGMA Toolbox* [72] to the Schaefer’s 400 brain regions [73] to explicitly visualize neural oscillations under different coupling strengths in Figure. 6**E**, and the results show that a high coupling value causes large neural synchronization.

**Figure 6.**
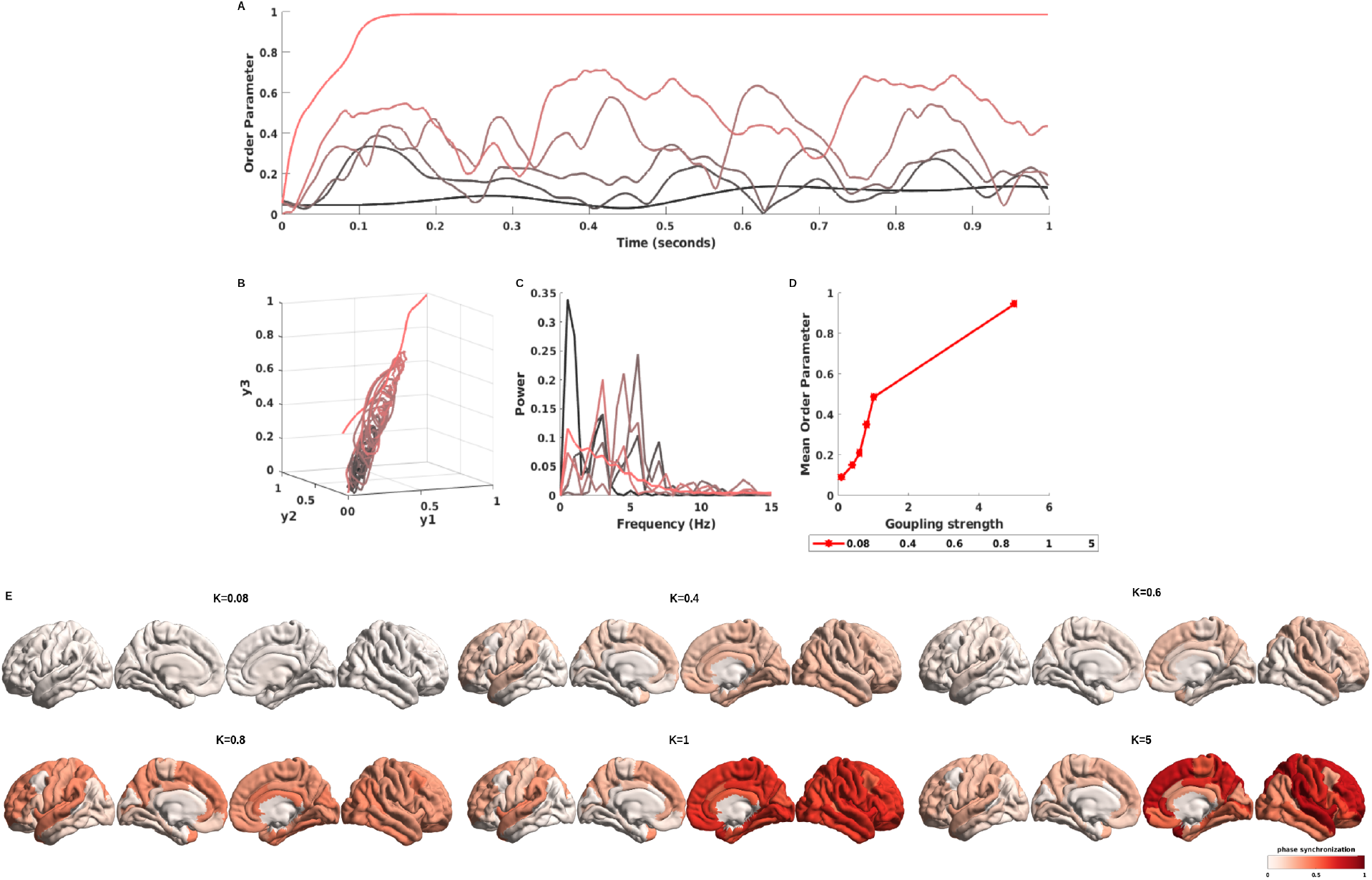
The order parameter of simulated BOLD signals with different coupling strengths. The **A** shows the order parameter of simulated BOLD signals with various coupling strengths (from weak coupling strength to high coupling strength, corresponding to dark black lines to red lines) from the Kuramoto model, and **B** shows the phase portrait of order parameter with simulated BOLD singals with *m* = 3 and *τ* = 1. The **C** shows power spectrum corresponding to order parameter of synthetic neural time series. The **D** shows the mean order parameter under different coupling strengths ([0.08, 0.4, 0.6, 0.8, 1, 5]). The **E** represented the mapping of phase synchrony to the brain surface and explained how the neural signal synchronized with a varying coupling strength value.

As previously stated, neural information interaction in the brain is constrained by structural connectivity, and moderate coupling between structural and functional connectivity drives collective neural activity, which then supports our high-level cognitive functions and behaviors. On the other hand, mathematically and biophysically realistic neural mass models can help us model the human brain functions, leading to a greater understanding of the fundamental cognitive functions of the brain and a better understanding of certain brain diseases. In this study, the FRP are used as functional connectivity descriptors to reconstruct predicted functional connectivity based on the Kuramoto model with a time delay.

The correlation between empirical functional connectivity, which includes 400 brain parcellation regions, and predicted functional connectivity from the mean Pearson correlation of FRP and the mean mutual information of FRP is reported in Figure. 7**B**, and **C**, respectively, after optimizing the model and searching for a range of free parameters in the model. The modest correlation values of *ρ* = 0.28 and 0.29 indicate that the Kuramoto model’s predicted functional connectivity based on FRP is performing well, and the correlation can be improved further by optimizing the free parameters in models and FRP.

**Figure 7.**
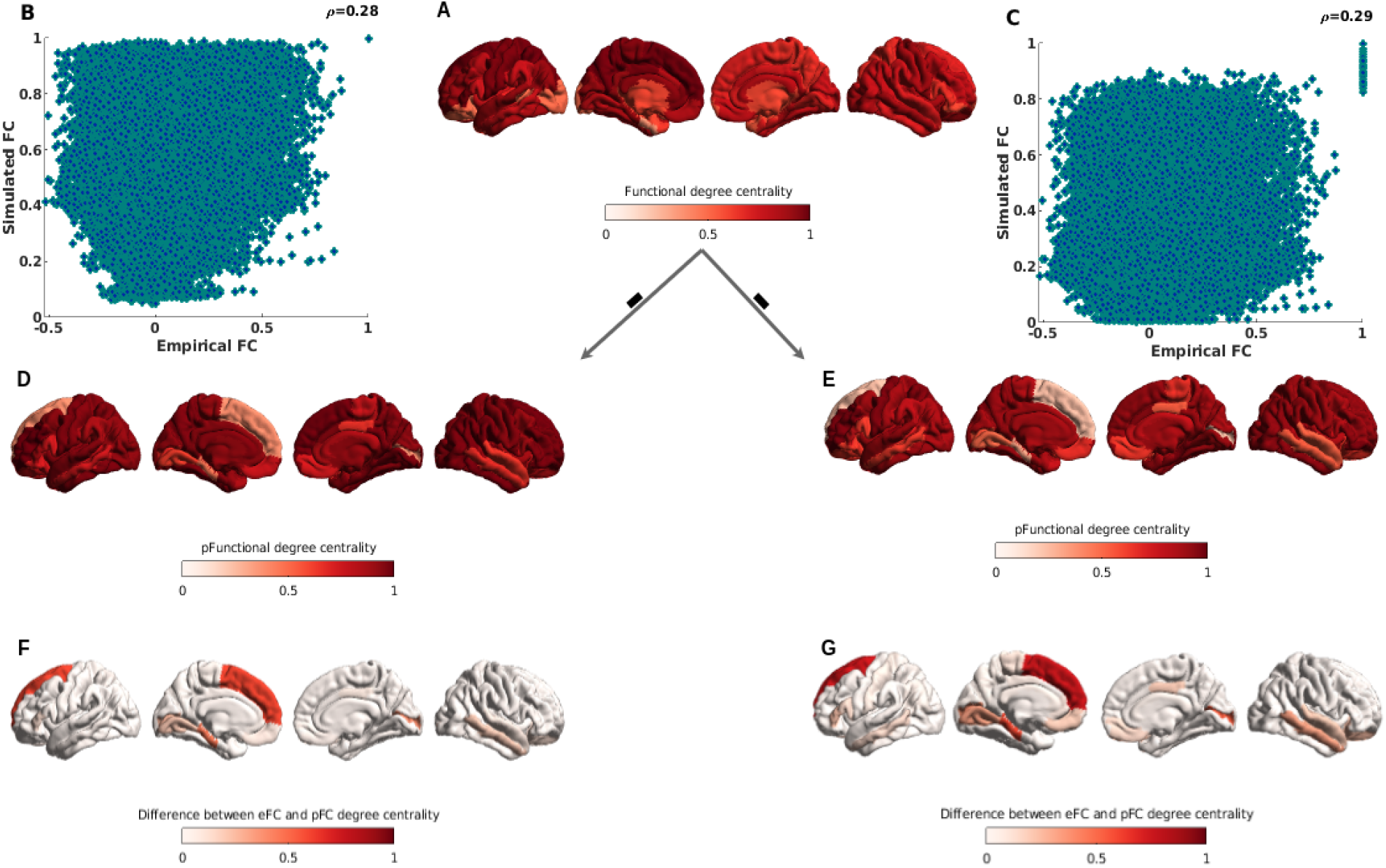
Cortical hubs of empirical functional connectivity, and predicted functional connectivity. The degree centrality used to identify functional hub regions. **A**.shows the functional degree centrality with empirical functional connectivity. **B**. and **C**. show the correlation between empirical functional connectivity and the mean Pearson correlation of predicted functional connectivity from FRP and the mutual information of predicted functional connectivity from FRP. **D**. and **E**. show the predicted functional degree centrality from predicted functional connectivity with mean Pearson correlation and mean mutual information of FRP. **F**. and **G**. show the topological difference between empirical and predicted functional degree centrality from correlation of FRP, and mutual information of FRP. See the main text for more information.

When it comes to relating macroscopic brain network organization, the structural and functional connectomes offer a wealth of useful information, and the brain networks appear to have few and well-localized regions with high functional connectivity density for fast integration of neural processing [74]. The hubs typically refer to regions of the brain with highly interconnected networks and richer connections. Using the aforementioned operations, we can determine the total weighted cortico-cortical connections for each region to generate weighted degree centrality maps, with higher degree centrality indicating hub regions. The degree centrality of empirical functional connectivity is measured and mapped to the brain surface in Figure. 7**A**, with darker red indicating more rich hubs. Meanwhile, we calculate the degree of centrality from predicted functional connectivity with FRP using Pearson correlation and mutual information, respectively, and then map both to the brain surface (Figure. 7**D**, and **E**), and to better discover the difference between empirical degree centrality and predicted degree centrality, the difference derived from empirical degree centrality minus predicted degree centrality, and from Figure. 7**F**, and **G**, we show that the biggest difference hubs are located in regions of the prefrontal cortex and in the ventral inferior temporal cortex, it explains that predicted functional connectivity from FRP has failed to build the right connectivity in the prefrontal cortex and ventral inferior temporal cortex, and both of these brain regions are responsive to high-level complex cognitive functions. However, the reasons may also be related to the neural mass model and related free parameters in both models and FRP, and making it a more proper and easy way to optimize these parameters will improve the accuracy of predicted functional connectivity. More details on this will be explained further in the discussion section. In general, our proposed approach successfully predicts the functional connectivity of the brain.

#### 2.4 Nonlinear Dynamics of the Functional Connectome in Brain Developmental fMRI

Here, we applied FRP to real fMRI data and aimed to use it as a descriptor of functional connectivity. In order to construct the complex brain networks, the whole data analysis pipeline is presented in Figure. 8. It provides an illustration of the schematic representation of the construction of brain networks using fMRI. After the time series have been extracted from the fMRI data using a functional atlas as a guide, the functional connectivity is estimated using phase portrait and FRP. The results are shown in a graph that shows both the brain’s nodes and how the functional connections between them are shown by the edge weights.

**Figure 8.**
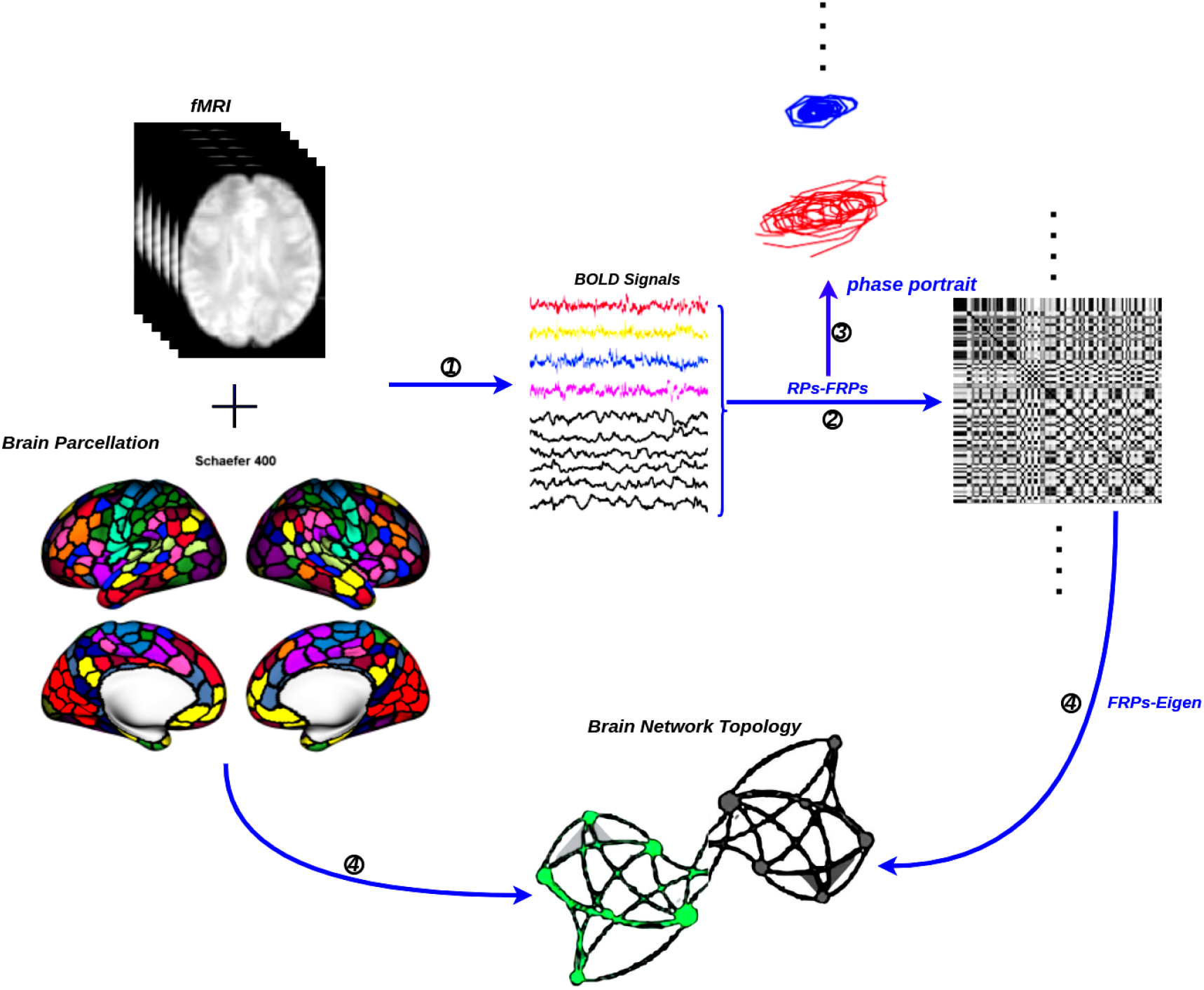
The general flowchart for the construction of a functional brain network by fMRI. ➀ Time series extraction from fMRI data within each functional unit (i.e., Schaefer’s 400 cortical regions [73]). ➁ Estimation of a functional connectivity with FRP. ➂ Examined the nonlinear properties of time series in the phase plane. ➃ Visualization of functional connectivity as a graphs network (i.e., network edges-FRP eigenvalue and network nodes-functional unit).

#### 2.4.1 Brain Developmental fMRI Dataset

We investigate the nonlinear dynamics behavior in fMRI data from brain developmental. The data mining efforts used open-source fMRI brain developmental studies involving 122 children (ages 3-12) and 33 adults (ages 18-39). In our study, we examined the fMRI scans of 66 people from the free and open-source *Nilearn* brain developmental dataset [75]. A panel of 33 children (aged 3 to 7) and 33 adults were surveyed (ages:18-39). While lying in the scanner, the participants watched Disney Pixar’s “Partly Cloudy” [76] without any additional tasks attached. After a long slumber, the movie began (black screen; TRs 0-5). This portion of the film is referred to as the “credits” (TRs 6-10). The functional data preprocessing details are described at https://osf.io/wjtyq/. The preprocessing neural signal can be directly downloaded from the *Nilearn* dataset^2^.

#### 2.4.2 The Pre-defined Functional Brain Atlas

We extracted BOLD signals from pre-defined brain atlas-MSDL [9], a brain atlas from resting-state fMRI scans multi-subject adaptation of sparse dictionary learning and estimate a group-level map of brain networks (see Figure. 9).

**Figure 9.**
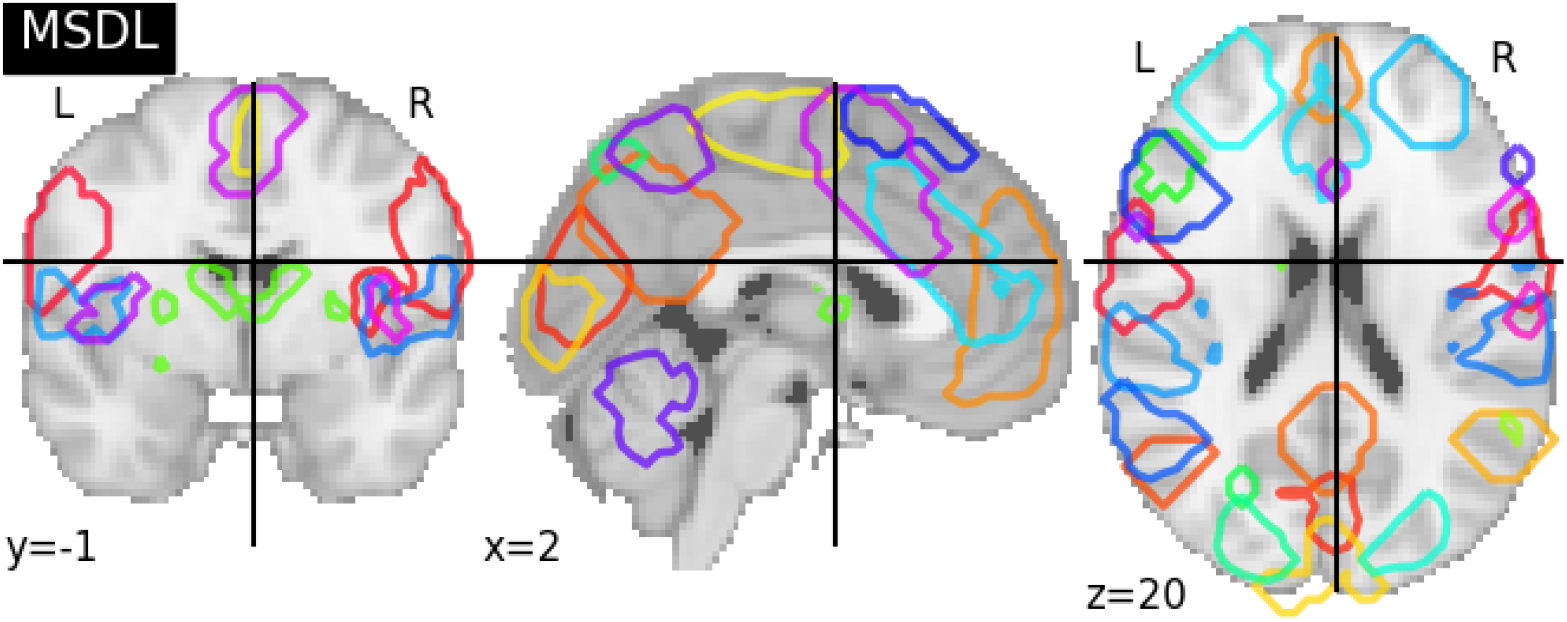
Pre-defined functional brain atlas-MSDL. The graph showed the volume of AAL (39 regions). The different brain areas are labeled on the brain volume with different octagon colors, and detailed ROI information can be found in the Appendix section with Table 4.

#### 2.4.3 Time Series Signals Extraction

The MSDL is a signal extraction method that combines a defining brain atlas with a bold signal extraction strategy. Averaging the voxels within each ROI, we extracted time series from each subject. Before moving on to the next step, we normalize and detrend time series.

#### 2.4.4 Nonlinear Dynamics of Brain Developmental of Neural Signals

As shown in Figure. 10, the phase portrait is first examined using the BOLD signals, and we displayed two examples from the L Aud and R LOC brain regions; the red line depicts an extracted neural signal from children, and the blue line represents a neural time series from adults; their related FRP and phase trajectories are estimated, and we can clearly see that both of the nonlinear dynamics descriptors capture different features.

**Figure 10.**
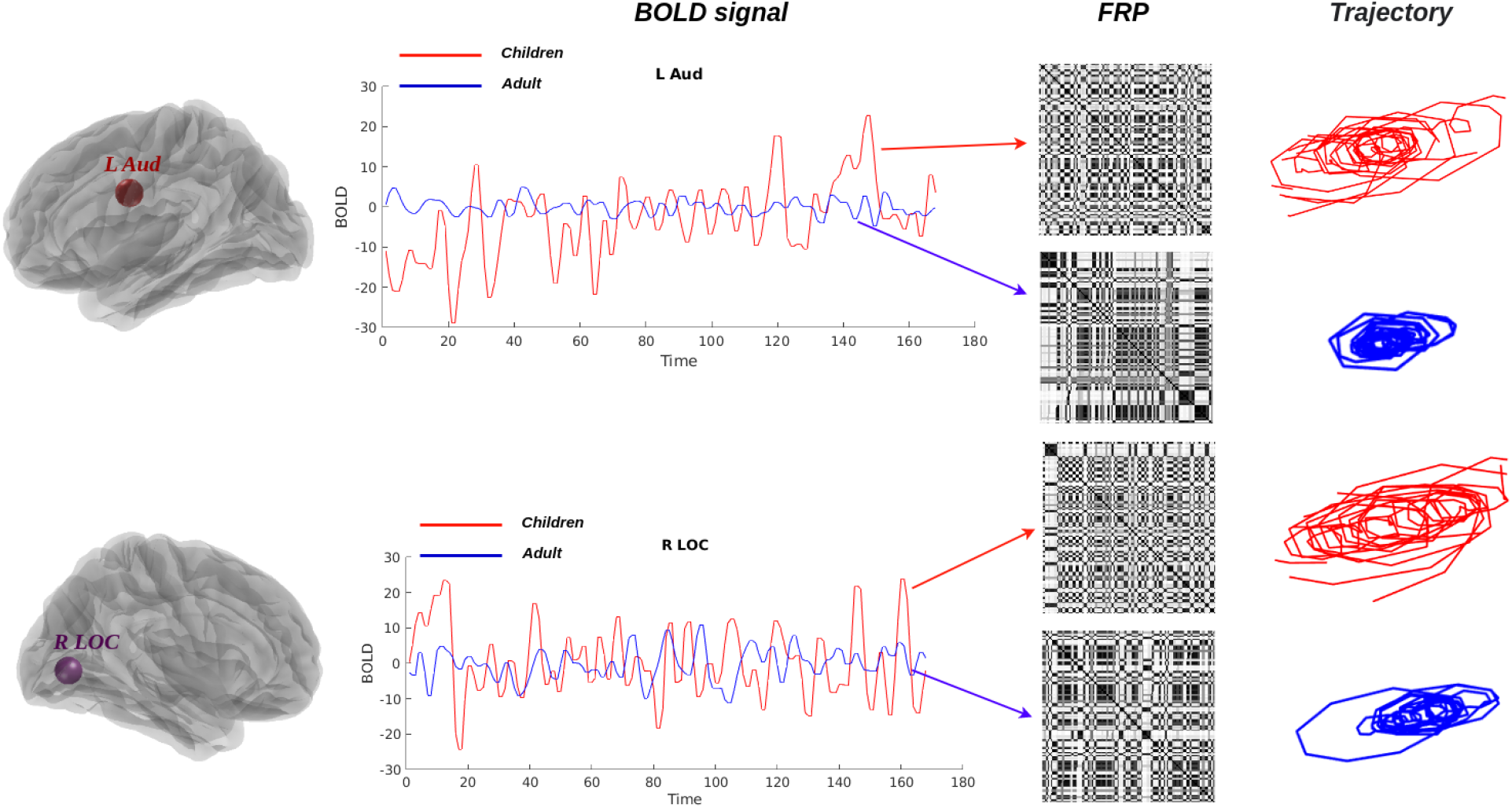
Brain fingerprinting with FRP and phase portrait. The variability of real neural BOLD signals (left panel) can be detected by FRP (middle panel) and related phase portraits (right panel). The red and blue lines depict the extracted BOLD signal from children and adults with two different brain regions and their variability, which exhibits a completely distinct pattern depending on the time interval.

In Figure. 11, we show the phase portrait result from one subject, both a child (red color) and an adult (blue color), from 39 brain regions. First, we discover that neural synchronization in most brain regions of the child is stronger than that of the adult under cartoon stimuli, and we suggest that can be a potential signature for explaining the reasons why children are more likely to watch cartoons than adults. Furthermore, the results also suggest that functional specialization increases throughout childhood, which is consistent with the original study [75]. However, the neural signals from children and adults still indeed share some similarities; for instance, all phase portraits are presented in limit-cycle-like patterns. Second, we suggest that the FRP can be used as brain fingerprinting metrics to capture the temporal pattern of neural signals, which indeed they do, as shown in Figure. 12, we estimate neural patterns from varying brain regions. Third, as we stated above, the features captured via phase portraits and FRP both share some similarities at some level, and we use a similarity index derived from a hierarchical clustering estimated from FRP to quantify the similarity, which is described in the following sections.

**Figure 11.**
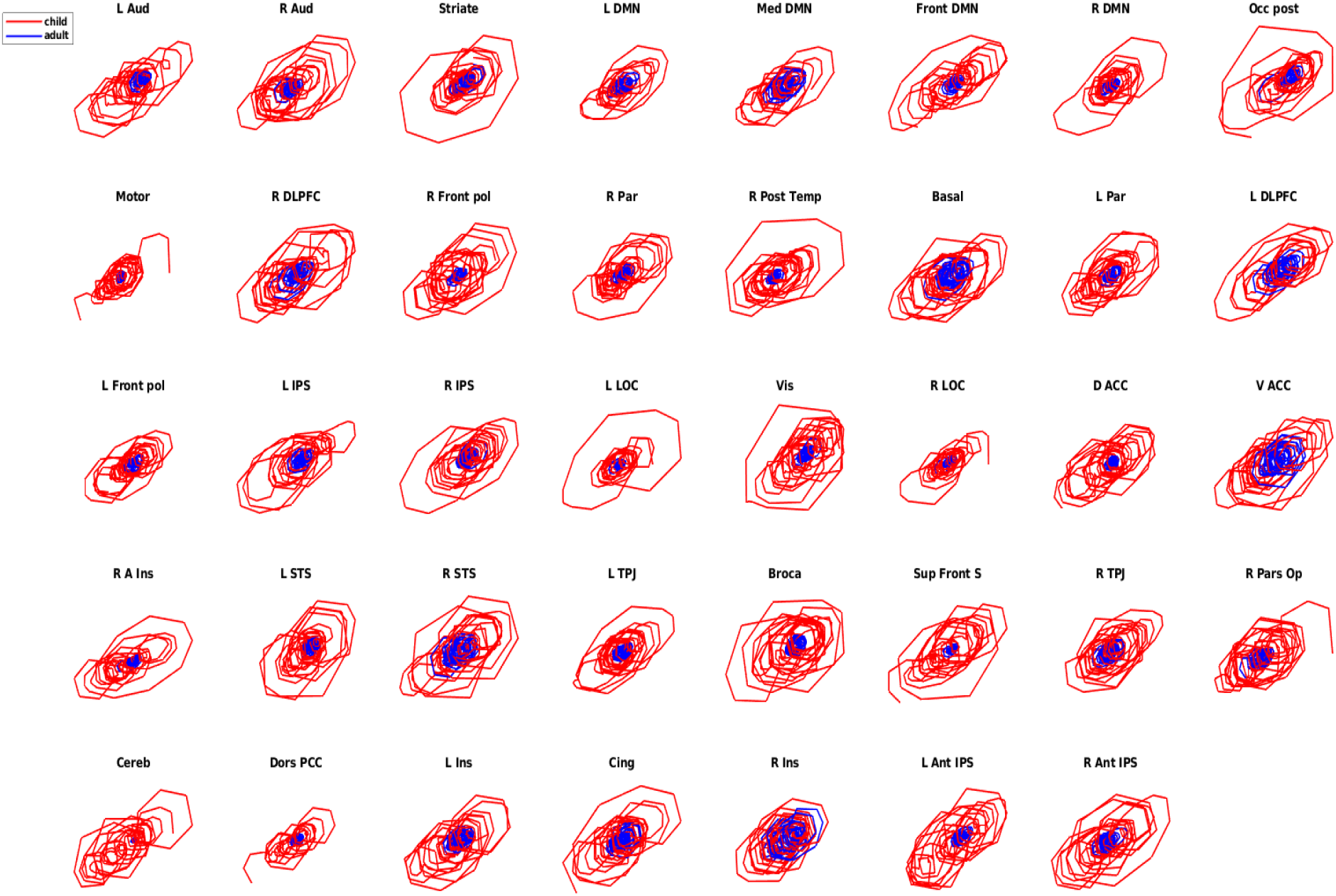
The phase portrait is a signature of the nonlinear dynamics of neural signals. The dynamics of real neural BOLD signals can be detected by their phase trajectories. The red and blue lines depict the phase portrait in children and adults from different brain regions, which exhibits a completely distinct pattern in children and adults.

**Figure 12.**
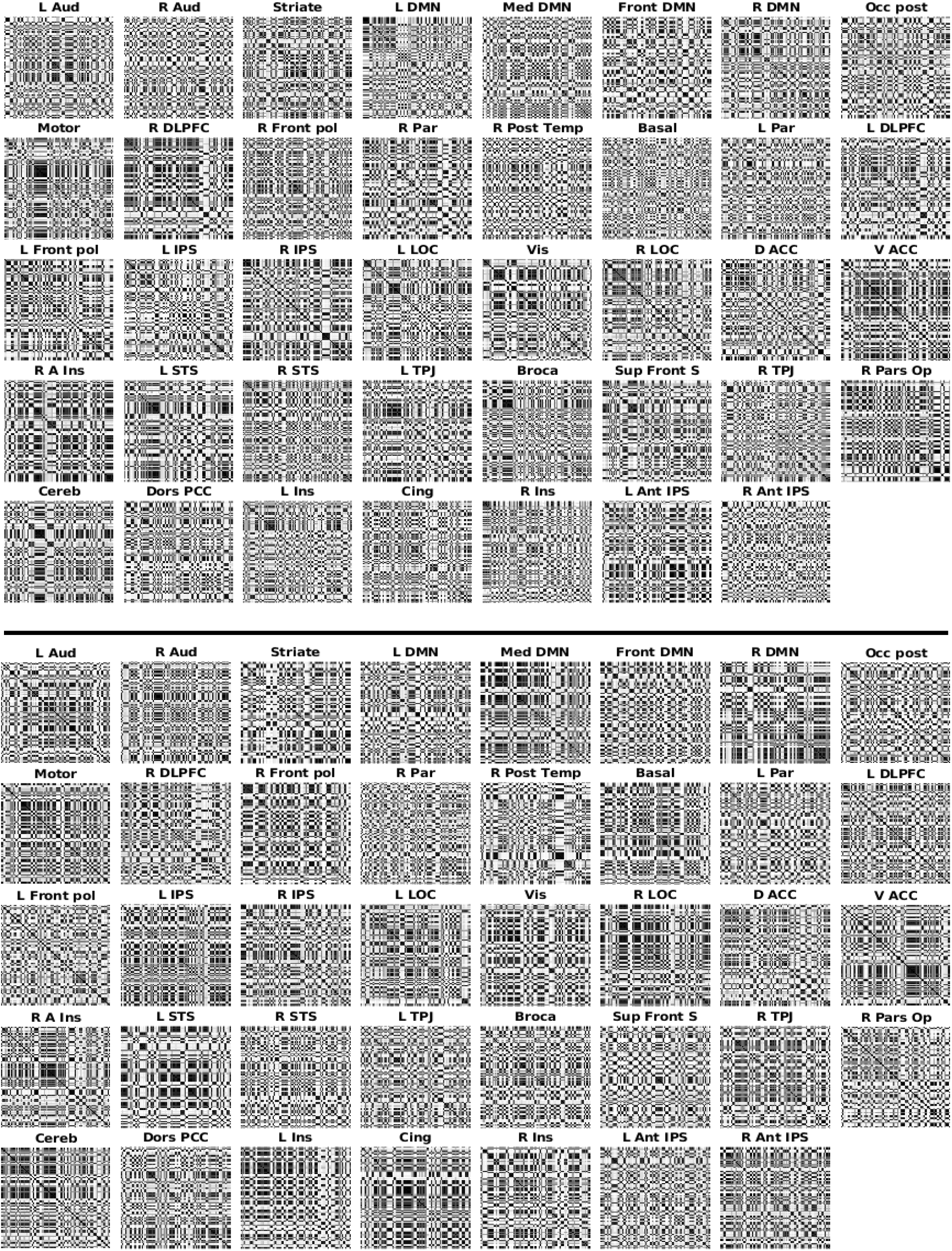
FRP are used to estimate functional connectivity. The figure depicts the BOLD signal from FRP in various brain regions, where the FRP were constructed with *m* = 3, *τ* = 1, *c* = 3. It can be seen that the texture of FRP in each brain region is different from other brain regions. The top plane shows the results from children, and the bottom plane represents the results from adults.

#### 2.4.5 Hierarchical Clustering Comparison

The discovery of hierarchical clusters in a dataset is the goal of the hierarchical clustering algorithm. Objects are initially grouped into a single cluster with the help of the clustering algorithm, which then divides the cluster into smaller sub-clusters based on their similarities. Comparisons of two objects’ similarities are typically calculated using distance metrics like the Euclidean distance. The algorithm keeps going until it reaches a point where there is no more division to be made. Here, we will use an index to quantify two hierarchical clusters [77].

Here we have two hierarchical clustering of the same number of objects, *n*. Let us consider the *N* = *n*(*n* − 1)*/*2 pairs of objects and let us define, for each non trivial partition in *k* groups,

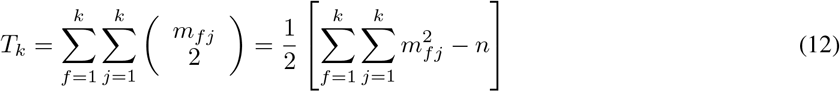

where *M*_*k*_ = [*m*_*fj*_] is a matching function, and *m*_*fj*_ indicates the number of objects placed in cluster *f* (*f* = 1, …, *k*), according to the first partition and in cluster *j*(*j* = 1, …, *k*). The Pair-joining counts in each partition are as follows,

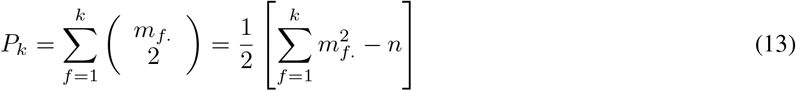

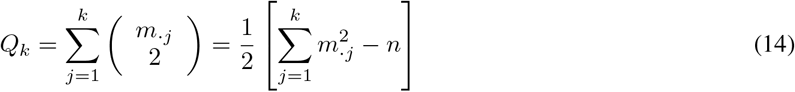

The index suggested by Fowlkes and Mallows [78] for two partitions in *k* groups is given by:

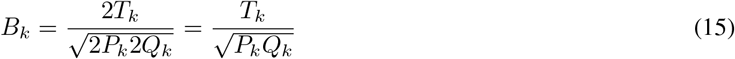

The similarity index *B*_*k*_ bounded in [0, 1]. *B*_*k*_ = 1 perfect matching between the two partitions, otherwise, *B*_*k*_ = 0 no matching between the two partitions.

Here, following this study [79, 80], we firstly feed the FRP features into a cascaded feature postprocessing that primarily uses convolution, rectification, and pooling components. Using an iterative procedure, the highest eigenvalue can be identified. Then we ran an agglomerative clustering (maximum clusters = 2) on the eigenvalue associated with 39 brain regions in all participants to see if the computed eigenvalue from FRP was associated with brain developmental. The corresponding hierarchical clustering result is presented in Figure. 13. We are able to quantify the differences in the brain developmental networks that children and adults go through as a result of aging by using hierarchical clustering to compare them. We quantify the similarity between adult and child brain networks under tasks, and the results suggest that adults and children share approximately *B*_*k*_ = 48.65% of their similarity on the similarity index. Therefore, the child and adult brain networks share some similarities, but they also have different brain network functions throughout the life span.

**Figure 13.**
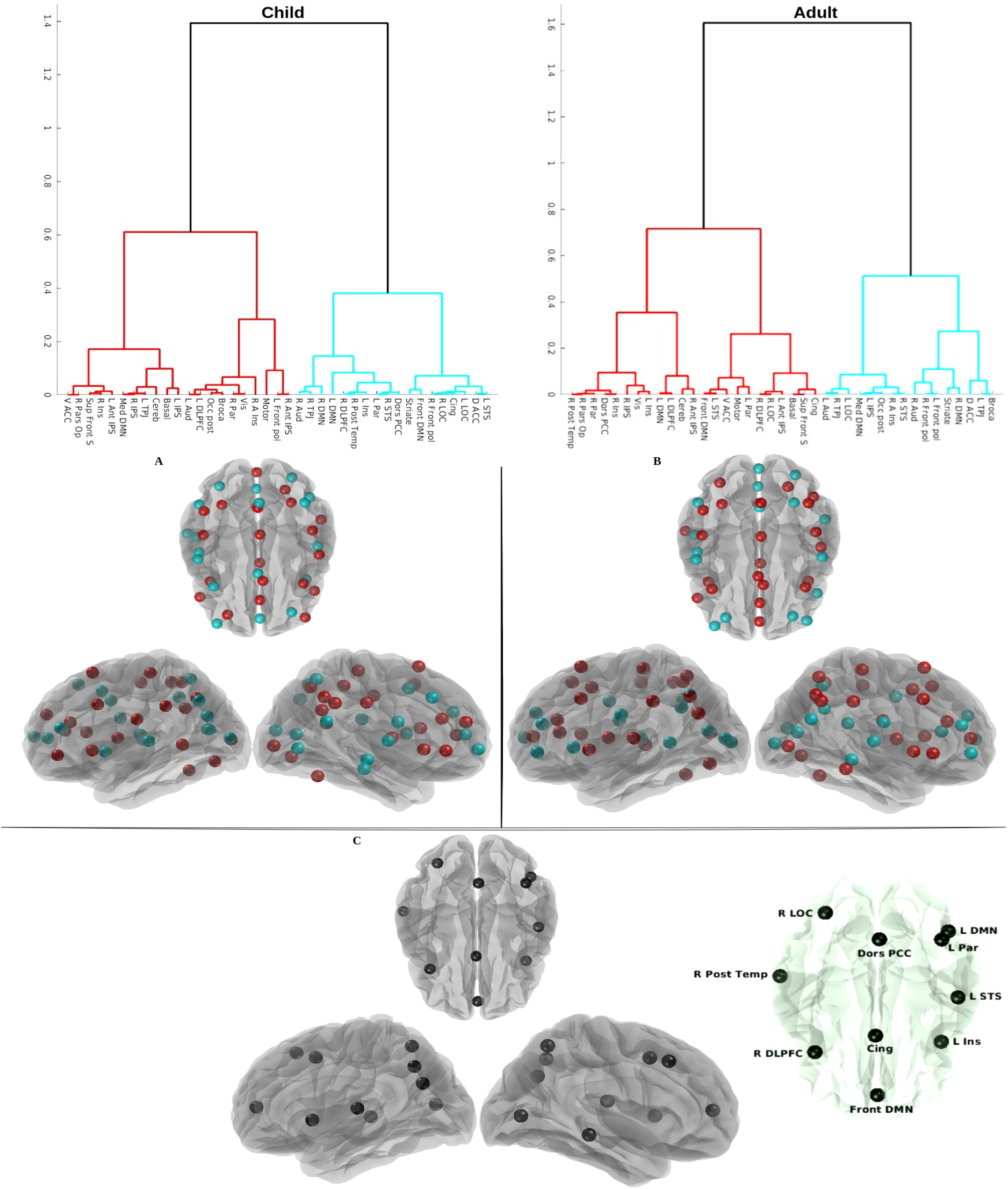
Hierarchical clustering for children and adults based on fuzzy recurrence eigenvalues. The top row left figure shows the hierarchical clustering result with children, and the top row right figure shows the clustering result with adults. The clustered brain areas are shown on the surface of the brain based on the results of the clustering in the top row, and colors (red and cyan) represent two groups of clustering branches for children (A) and adults (B) in the middle row, respectively. The topological brain clustering difference between children and adults is shown in the last row (C), and the difference in brain regions is shown with black nodes on the brain surface, such as R LOC, R Post Temp, R DLPFC, Front DMN, Cing, Dors PCC, L DMN, L Par, L STS, and L Ins. The detailed explanations were presented in the main text.

In order to better visualize the relationship between clustering brain regions, we projected the two classes (red and cyan) of clustering brain regions in children(Figure. 13.**A**) and adults(Figure. 13.**B**) on the surface of the brain. The topological difference was plotted in Figure. 13.**C**. The main differences exist in visual areas, attentional response regions, default model networks, and temporal response regions, and the listed brain regions, such as R LOC, R Post Temp, R DLPFC, Front DMN, Cing, Dors PCC, L DMN, L Par, L STS, and L Ins may play an important cognitive role in the development of the Theory of Mind^3^, specifically from early life to adulthood, which is consistent with previous findings from human behavior experiments [81, 82].

## 3 Discussion

Our research expands our understanding of nonlinear dynamics in human complex brain networks in four key directions, with implications for biophysical theory and practice. First, we presented novel methods based on phase portraits and FRP for investigating the nonlinear dynamics behaviors of neural signals. We found that the method can, in fact, capture the nonlinear dynamics of neural time series. The phase trajectories forming a flow on the manifold and the sets of these flow curves forming orbits, as well as the phase trajectories of the generated neural signals forming an attractor, tend to enter limit cycles with increasing time in our numerical experiments, which include physically realistic and bio-physically realistic synthetic neural signals. The same phenomenon also happened in the real BOLD signals, which explains the chaotic behavior of neural activity. Furthermore, the presence of low-dimensional attractors in both numerical and real-world experiments supports the involvement of chaos as a fundamental aspect of brain dynamics, and its consistency and ability to explain chaotic dynamics in the human brain may be of functional utility in central nervous system cognitive processes [70, 83]. Moreover, phase portraits and FRP make brain fingerprinting are possible because we see that both of them can capture the small variability of neural signals with time change.

Second, we investigated the limited examples of nonlinear dynamics in models and examined their nonlinear dynamics behaviors in numerical experiments. However, as we mentioned, we did not explore too many nonlinear dynamics models, and we should try more nonlinear dynamics models, including both physics- and biophysics realistic neural mass models, which will explain different dynamics behaviors. Further, there are some certain free parameters in dynamics models, and their parameters directly affect the nonlinear dynamics behaviors in the model, and we have not fully explored these parameters. The most important case is the neural mass model in our simulation case; here, we only use it to emphasize that the FRP can predict the functional connectivity based on structure connectivity, and we have not explained too much the significant meaning between the connections between structure and functional connectivity. Furthermore, the correlation between empirical functional connectivity and predicted functional connectivity can be improved with more appropriate ways to optimize the free parameters in models, and we still need to consider other ways to measure the spatial-temporal properties of each brain region FRP, while not ignoring the high-order connectivity patterns.

Lastly, in the real experiment, we measured the brain networks from FRP-eigenvalues via hierarchical clustering, then the similarity index between both brain networks was investigated in children and adults. we discovered approximately 48.65% similarity between the two groups. This finding suggests that brain development is still ongoing and that children’s brains are not yet fully developed, and the neural architecture underlying cognitive processes continues to mature throughout childhood and adolescence. The brain undergoes significant changes during these stages. Meanwhile, we also identified specific regions with notable differences, including the DLPFC, PCC, DMN, and STS. These brain regions play a crucial role in constructing social cognitive processing networks in the human brain, which are primarily related to social tasks [81, 82]. However, in this analysis, we did not consider high-order brain connectivity information and only captured pairwise relationships. Therefore, we need to explore and apply alternative methods to measure brain network connectivity that retain both low-order and high-order connectivity patterns [35–39].

Furthermore, in the empirical experiments, we used the open source data to explore the dynamics of neural activity, and it will be interesting for us to apply these metrics to a dataset of cognitive tasks or a dataset of brain diseases, and we are sure that will capture some interesting patterns to help us explain some brain functions or find some biomarkers for brain diseases [39]. In the future, it will be our branch extension work. Moreover, in order to make the result reliable, we should consider applying it to a large fMRI dataset and make sure the capture pattern of brain networks is stable and can be reproduced.

Taken together, the first thing we discovered through our numerical experiments was that FRP was very sensitive to capture the changes in neural signals, that the associated nonlinear dynamic behavior can be described by a temporal trajectory in phase space, and that by examining the nonlinear dynamics of complex brain networks, we can also capture the regular and irregular brain networks and then understand the brain dynamics from there. Second, we discovered that the FRP may represent the fingerprinting functional connectivity of the brain regions from adjacency and long-distance brain areas when using fMRI, which makes brain fingerprinting possible. Finally, through our studies, we suggested that phase trajectories and FRP could be highly effective functional connectivity descriptors for explaining brain dynamics, and they have great potential when applied to clinic applications.

## 4 Conclusions

Through numerical studies, we found that FRP were highly sensitive to variations in neural signals, and the resulting nonlinear dynamic behavior could be characterized by a temporal trajectory in phase space. This was a significant finding, as it clarified that the nonlinear dynamics of neural signals could be captured using phase portraits and FRP. Second, we found that the functional variability of neural signals, as observed through fMRI, can be represented by FRP. The phase trajectory and FRP served as metrics to explain the nonlinear dynamics of brain networks. Third, we analyzed a brain developmental fMRI dataset using fuzzy recurrence eigenvalues to group brain development into visually understandable units that illustrated distinct brain regions associated with aging. By using FRP eigenvalues, we constructed meaningful biological functional connectivity, leading to robust and rich latent information that enhanced our understanding of lifespan development. Lastly, we concluded that, in the context of nonlinear dynamics, phase trajectories and FRP can serve as particularly useful descriptors of functional connectivity.

## Data Availability

Human neuroimaging data used in this study were provided by the Human Connectome Project (HCP) [12] (https://www.humanconnectome.org), WU-Minn Consortium (Principal Investigators: David Van Essen and Kamil Ugurbil; 1U54MH091657) funded by the 16 NIH Institutes and Centers that support the NIH Blueprint for Neuroscience Research; and by the McDonnell Center for Systems Neuroscience at Washington University. The normative connectomes were computed from Human Connectome Project data and included as part of the local structure-function toolbox [69] (https://github.com/brain-networks/local_scfc). Simulated data are available from the corresponding author on reasonable request. The open-source brain developmental fMRI data collected and analyzed during the current study are available through the OpenfMRI project (https://openfmri.org/;Link:https://www.openfmri.org/dataset/ds000228/DOI:10.5072/FK2V69GD88).

## Acknowledgement

Data were provided, in part, by the Human Connectome Project, WU-Minn Con sortium (principal investigators: D. Van Essen and K. Ugurbil; 1U54MH091657), funded by the 16 NIH Institutes and centers that support the NIH Blueprint for Neuroscience Research and by the McDonnell Center for Systems Neuroscience at Washington University.

The authors would like to acknowledge the use of the following freely available code:

**Ghost Attractors**, https://github.com/jvohryzek/GhostAttractors.

**Hopf Delay Toolbox**, https://github.com/fcast7/Hopf_Delay_Toolbox.

**The ENIGMA TOOLBOX**, https://github.com/MICA-MNI/ENIGMA.

**Local Structure-Function**, https://github.com/brain-networks/local_scfc.

**Fuzzy Recurrence Plots**, https://drive.google.com/file/d/189HUUNZGSjdsql3lmGI8oXdTeUrRCps0/view.

## Ethics Statement

All human data used in this study is from the public repository, which is distributed in compliance with international ethical guidelines.

## Appendix

**Table 4.**
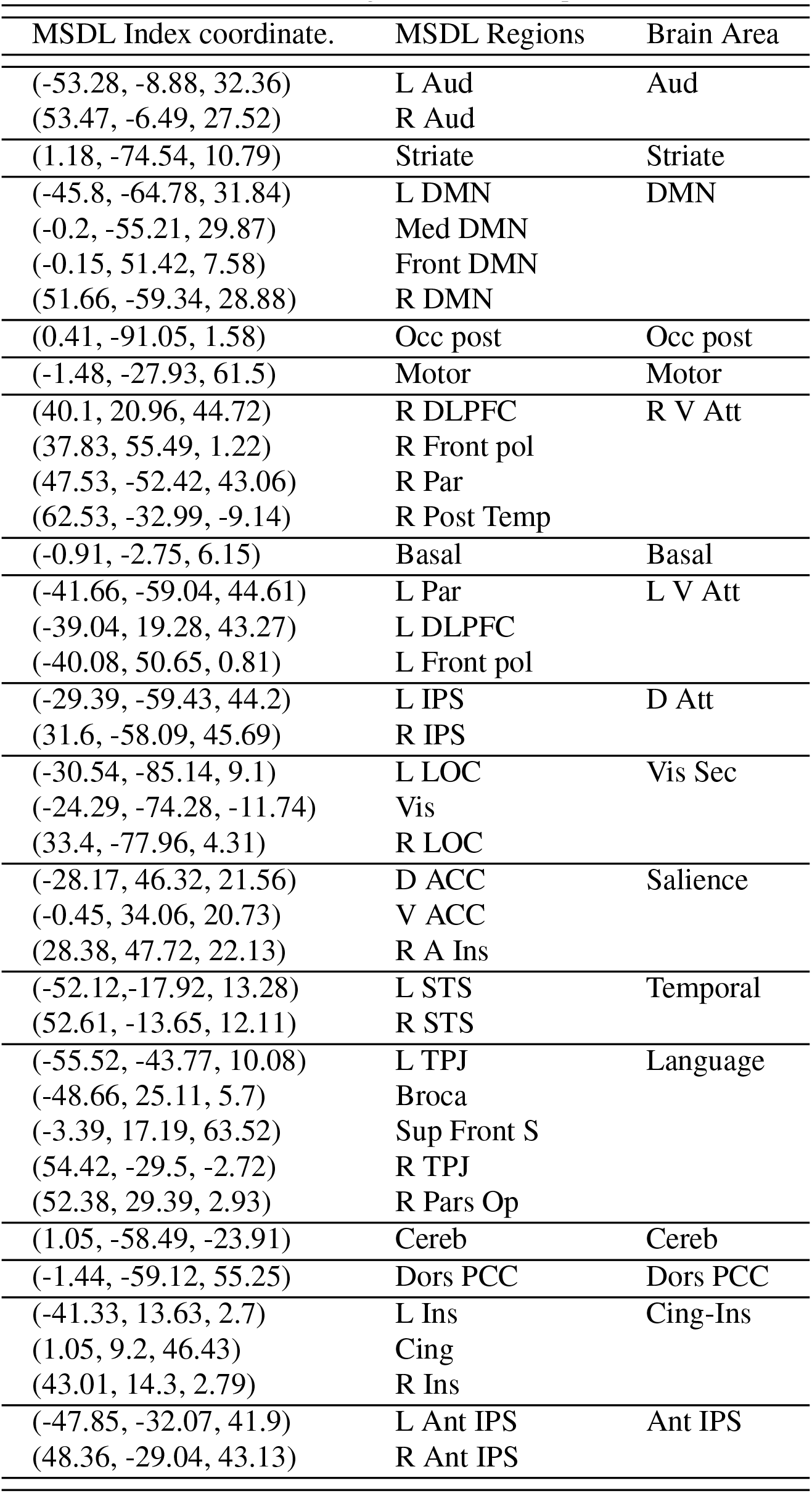
Information of 39 brain regions that are comprised in the MSDL atlas.

## Footnotes

1 http://www.recurrence-plot.tk/index.php

2 https://nilearn.github.io/dev/modules/generated/nilearn.datasets.fetch_development_fmri.html

3 Theory of Mind is the ability to attribute mental states to ourselves and others, serving as one of the foundational elements for social interaction.

